# White matter in infancy is prospectively associated with language outcome in kindergarten

**DOI:** 10.1101/781914

**Authors:** Jennifer Zuk, Xi Yu, Joseph Sanfilippo, Michael Joseph Figuccio, Jade Dunstan, Clarisa Carruthers, Georgios Sideridis, Borjan Gagoski, Patricia Ellen Grant, Nadine Gaab

## Abstract

Language acquisition is of central importance to a child’s development. Although the trajectory of acquisition is shaped by input and experience postnatally, the neural basis for language emerges prenatally. Thus a fundamental question remains unexamined: to what extent may the structural foundations for language established in infancy predict long-term language abilities? In this longitudinal neuroimaging investigation of children from infancy to kindergarten, we find that white matter organization in infancy is prospectively associated with subsequent language abilities, specifically between: (i) the left arcuate fasciculus in infancy and subsequent phonological awareness and vocabulary knowledge, and (ii) the left corticospinal tract in infancy and phonological awareness and phonological memory in kindergarten. Results are independent of age and home literacy environment. These findings directly link white matter organization in infancy with language abilities after school entry, and suggest that structural organization in infancy sets an important foundation for subsequent language development.

## INTRODUCTION

The neural basis for language has been postulated to emerge prenatally, suggesting that the rudimentary foundation for language has already been established in the brain by the time an infant is born (Dehaene-Lambertz and Spelke, 2015). Yet the complex trajectory of language acquisition extends well beyond childhood (Owens, 2008). Environmental input and experience over time play a crucial role in shaping the trajectory of language acquisition (Rowe, 2012; Weisleder and Fernald, 2013), but the first two years of life signify a particularly rapid period of language development and brain maturation (Pujol et al., 2006), and are increasingly recognized to set an important foundation for long-term development (Gilmore et al., 2018). To date, it remains unknown whether the neural scaffold for language established in infancy is prospectively associated with the protracted trajectory of language development. Here we provide the first longitudinal evidence of prospective associations between brain structure in infancy and language skills several years later, at the age when children start formal schooling.

While environmental exposure and experience are essential for language acquisition, biological determinants also make an integral contribution during prenatal and early postnatal development (Graham and Fisher, 2013). Structural organization of white matter is one critical component of neuroarchitecture that underlies efficient signal transmission and corresponding skill acquisition (including language), and undergoes a robust period of development within the first two years of life (Gilmore et al., 2018). Although the myelination of axons that comprise white matter increases throughout early childhood and beyond (Barnea-Goraly et al., 2005; Dubois et al., 2008; Lebel et al., 2008; Mukherjee et al., 2001), this process is characterized by reorganization and plasticity that builds upon the microstructure established during infancy (Gilmore et al., 2018). The most rapid production of myelin occurs during prenatal and early infant development (Lenroot and Giedd, 2006), in conjunction with dynamic changes in gene expression in brain tissue (Naumova et al., 2013). Studies of whole-genome expression in the developing brain reveal that the temporal dynamics of the transcriptome are more robust during prenatal development than at any postnatal stage (Colantuoni et al., 2011; Johnson et al., 2009; Kang et al., 2011; Lambert et al., 2011; Somel et al., 2010). Moreover, candidate susceptibility genes for speech and language disorders show an especially prominent window of expression during prenatal development in the perisylvian cortex, the part of the brain that encompasses the neural basis of language (Abrahams et al., 2007; Johnson et al., 2009). Thus, an important white matter foundation is established in infancy that sets children on a path for subsequent development.

Although the first two years of life mark a peak in the rate of white matter development, longitudinal studies tracking long-term language development from infancy have predominantly employed behavioral and electrophysiological methods. Behavioral and functional neural responses to tones and speech syllables in infancy are prospectively associated with the number of words a child understands and produces at the toddler stage (Friedrich and Friederici, 2006; Kuhl et al., 2008; Tsao et al., 2004), and subsequent phonological awareness and oral comprehension abilities at the preschool age (Benasich, 2002; Guttorm et al., 2010; Guttorm, 2005; Leppanen et al., 2010; Molfese and Molfese, 1997). These findings illuminate foundations for language in infancy that are predictive of subsequent language outcomes from a functional standpoint, yet what is the role of underlying white matter organization? Converging evidence supports the notion that white matter connectivity precedes function, as revealed by patterns of structural connectivity at preschool age shown to predict subsequent functional development (Saygin et al., 2016). Crucial for language, the arcuate fasciculus connects frontal and temporoparietal brain regions, which are responsible for language production and comprehension, respectively (Brauer et al., 2011; Catani et al., 2005; Skeide et al., 2016). This dorsal route is a key component of the superior longitudinal fasciculus, which together with ventral routes such as the inferior longitudinal fasciculus (a tract that connects anterior temporal and posterior occipitotemporal regions, Catani et al., 2003), comprise a comprehensive network of white matter that underlies the integration of language comprehension, formulation, and expression (Dick et al., 2014). Recent methodological advances to conduct structural neuroimaging in infancy reveal that these white matter pathways essential for language emerge within the first year of life (Brauer et al., 2013; Dubois et al., 2009; Dubois et al., 2016; Geng et al., 2012; Leroy et al., 2011). Although converging evidence suggests that language specificity within white matter is established in infancy, no study to date has examined how this relates to long-term language abilities.

Relationships between white matter and language are specific and independent of socioeconomic factors according to evidence acquired at the school age (Romeo et al., 2018), but have yet to be investigated starting from infancy. Among school-age children, structural organization of the left arcuate fasciculus has been positively linked with language skills in domains including phonological awareness and vocabulary knowledge (Lebel and Beaulieu, 2009), and cross-sectional links with phonological awareness have been shown as early as three years of age (Reynolds et al., 2019). This pathway has also been directly associated with language exposure (i.e., the number of conversational turns in dialogic interaction) independent of socioeconomic status (Romeo et al., 2018). In fact, structural organization of the left arcuate fasciculus was observed to mediate the well-known relationship between language exposure and language skills among four-to six-year-old children (Romeo et al., 2018). Therefore, language-related white matter organization has been indicated as a specific and crucial mechanism underlying language abilities; however, there has yet to be any indication of how early white matter in infancy may predict subsequent language abilities at the school age.

Here, we examine prospective longitudinal relationships between white matter organization in infancy and subsequent language abilities in kindergarten. The present longitudinal investigation initially acquired structural neuroimaging in infancy, then followed children until they reached the kindergarten age, at which time they completed a standardized battery of measures that characterize several aspects of language abilities. Our findings reveal selective prospective associations between the structural organization of white matter in infancy and language abilities in kindergarten that are robust even when accounting for age and factors related to the child’s home environment. These findings are the first to suggest that white matter within the first two years of life serves as an important foundation underlying the developmental trajectory of language development in early childhood.

## RESULTS

Forty children who completed diffusion-weighted imaging in infancy and also returned for longitudinal follow-up (4-5 years later) were included in the present analysis as part of a larger longitudinal investigation tracking brain and language/pre-literacy development from infancy to late elementary school (the BabyBOLD study). Neuroimaging was initially acquired with healthy, full-term infants (mean age at infancy: 10 months; age range: 4-18 months) while the infants were naturally sleeping (Raschle et al., 2012). Longitudinal follow-up at the school age (mean age: 5.5 years, age range: 4–6.5 years) involved behavioral standardized measures that characterize phonological skills, oral comprehension, and vocabulary knowledge. The longitudinal timeframe between infant and follow-up time points was 4.61 ± 0.65 years on average. We employed multiple regression analyses to examine relationships between white matter pathways in infancy and subsequent language abilities at the school age.

Models were constructed for each of the left-hemispheric white matter tracts of interest, with the left arcuate fasciculus as the primary pathway implicated in language and the left inferior longitudinal fasciculus as a ventral complement to this dorsal pathway. We also incorporated the left corticospinal tract to examine whether potential effects are specific to language-related tracts or may be reflected in basic motor connections subserving language (Walton et al., 2018). Age at the neuroimaging session in infancy constituted a control predictor to account for potential age effects in white matter organization, and standard scores for the language measures were defined as the outcome variables. Socioeconomic status was controlled by minimal variation in the present sample, but key factors characterizing the home literacy environment were included in the model (amount of time spent reading to the child, number of children’s books in the home), since shared parent-infant book reading has been linked with language acquisition in infancy (Karrass and Braungart-Rieker, 2005) and prospectively associated with subsequent language abilities (Raikes et al., 2006). Multiple regression models were run for each white matter tract with infant age and home literacy variables as predictors, and each subsequent language outcome at the school age. We employed parametric bootstrapping to simulate the sampling distribution in the general population in order to properly evaluate the magnitude of the present effects, and utilized Family-Wise Error (FWE) rate adjustment as the threshold of significance to correct for multiple comparisons with each model.

### Prospective associations between language-related white matter organization in infancy and subsequent language abilities

With the left arcuate fasciculus (AF) as our predictor, holding constant the effects of infant age and key factors related to the home literacy environment, fractional anisotropy (FA) within the posterior segments of this tract in infancy is prospectively associated with several of the language outcome measures. Specifically, FA in the posterior segment of the left AF in infancy is prospectively associated with subsequent phonological awareness abilities, as indicated by Segmentation and Sound Blending measures from the Woodcock Johnson Oral Language Assessment (*WJ-OL IV*, FWE-corrected, *p* < 0.005, Figure 1). On average across all significant nodes identified within the posterior segment of this tract in relation to Segmentation (nodes 38–50), infant-age predictors in the model explain 35% of the variance in kindergarten-age segmentation skills. For the significant nodes associated with Sound Blending (nodes 48–50), the model explains 28% of the variance in sound blending abilities. Significant prospective associations were also identified between FA in the posterior segment of the left AF in infancy and subsequent vocabulary knowledge (FWE-corrected, *p* < 0.006, nodes 48-50, see Figure 1), as measured by the Peabody Picture Vocabulary Test (*PPVT-IV*). Among the significant nodes implicated, the model explains 21% of the variance in kindergarten-age vocabulary knowledge.

**Figure 1 Legend:**
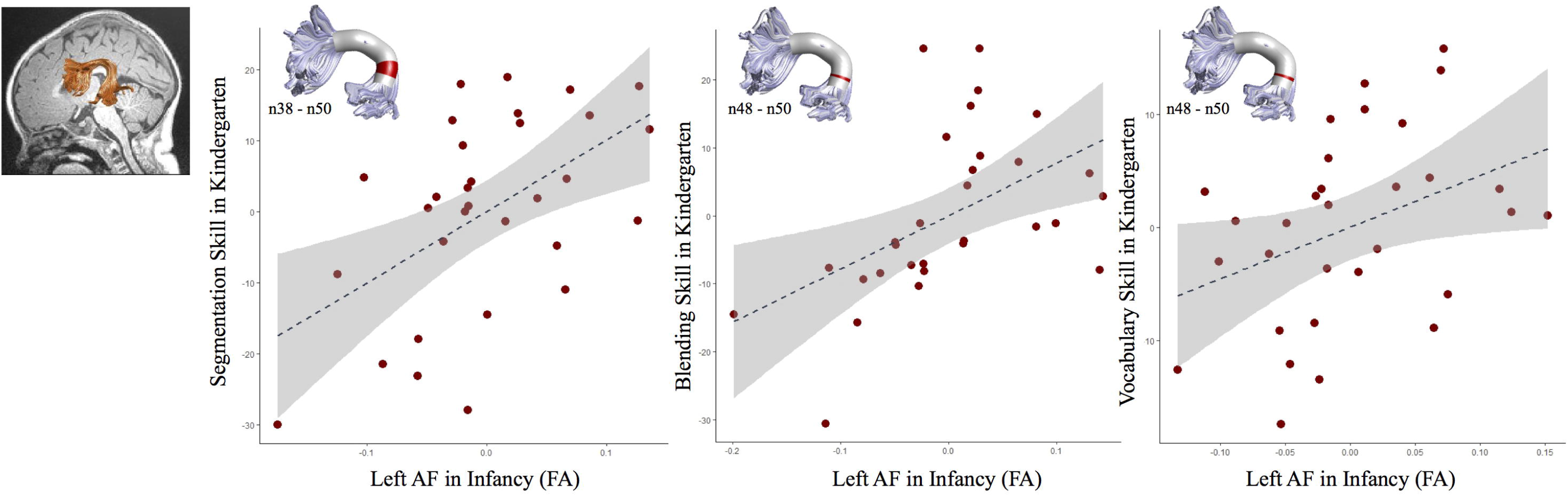
FA of the left arcuate fasciculus (AF) in infancy is prospectively associated with phonological awareness and vocabulary knowledge skills in kindergarten. Significant associations between FA of the posterior segment of the left AF in infancy and language skills in kindergarten (displayed in terms of the centered residuals produced from partial correlations). Nodes in the posterior segment of the left AF that show significant effects are marked in red on the 3-dimensional rendered tract (*p* < 0.05, FWE-corrected).

No prospective relationships were identified between FA of the inferior longitudinal fasciculus in infancy and kindergarten-age language abilities when holding constant the effects of infant age and key factors related to the home literacy environment.

### Corticospinal tract in infancy is prospectively associated with subsequent phonological abilities

To determine whether our data point towards language-specific or also more basic motor-based effects, we investigated the corticospinal tract as well. Here we find that the left corticospinal tract is prospectively associated with subsequent phonological skills at the school age. Holding constant the effects of infant age and key factors related to the home literacy environment, FA in nodes 1-12 of this tract are prospectively associated with phonological awareness (as indicated by Segmentation), and nodes 10-15 are associated with phonological working memory skills (as indicated by the Memory for Digits subtest from the Comprehensive Test of Phonological Processing, *CTOPP-2;* FWE-corrected, *p* < 0.007, see Figure 2). On average across all significant nodes identified within this tract for each outcome measure respectively, predictors were found to explain 42% of the variance in kindergarten-age segmentation skills, and 13% of the variance in phonological memory abilities.

**Figure 2 Legend:**
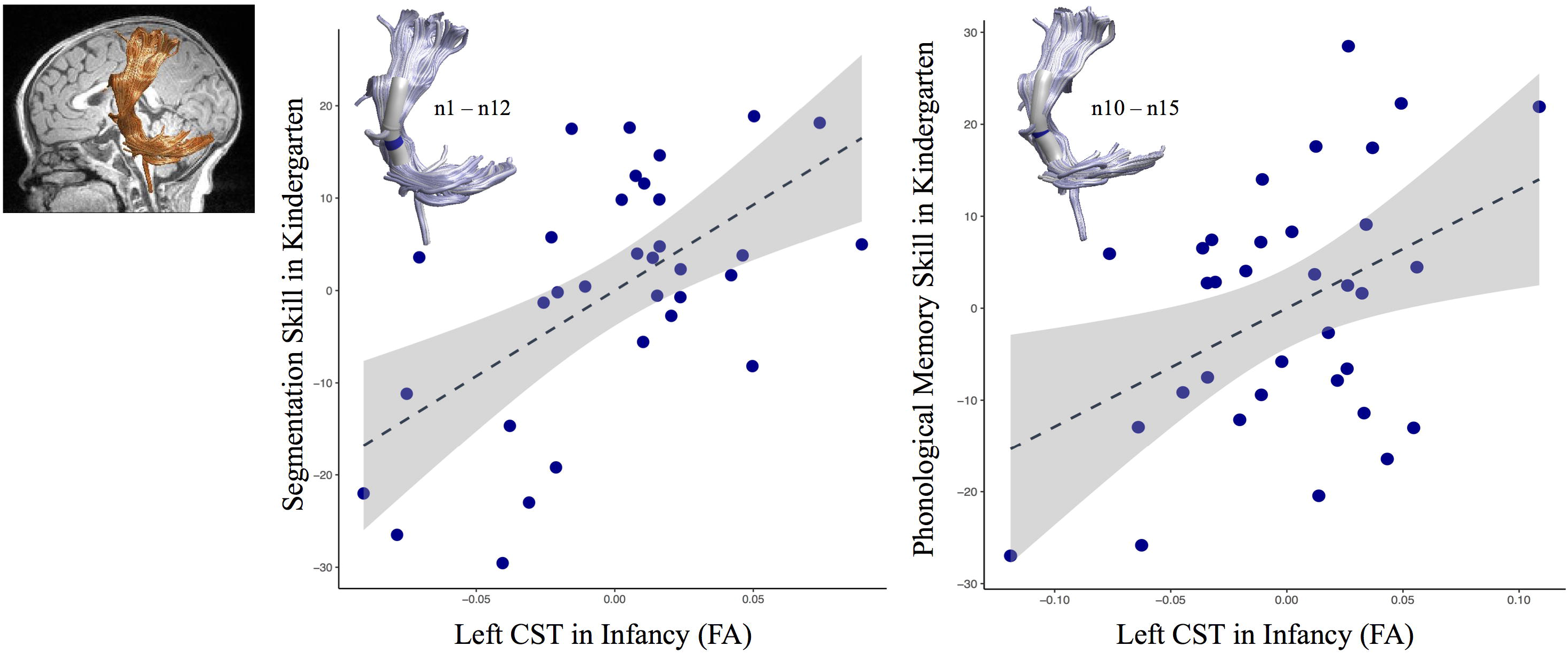
FA of the left corticospinal tract in infancy is prospectively associated with phonological awareness and phonological memory in kindergarten. Significant associations between FA of the left corticospinal tract in infancy and phonological skills in kindergarten (displayed in terms of the centered residuals produced from partial correlations). Nodes of the left corticospinal tract that show significant effects are marked in blue on the 3-dimensional rendered tract (*p* < 0.05, FWE-corrected).

### Contributions of the arcuate fasciculus and corticospinal tract in infancy to the prediction of kindergarten-age phonological awareness abilities

Based on our findings within the left AF and corticospinal tract in association with kindergarten-age phonological awareness abilities (as indicated by Segmentation), we built a final model incorporating both tracts while also accounting for infant age and key factors related to the home literacy environment. Incorporating the average FA across significant nodes identified in previous analysis models for each tract, both the left AF (β = .386, *p* < 0.05) and left corticospinal tract (β = .608, *p* < 0.005) are prospectively associated with kindergarten-age segmentation skills in this model. Overall, the model explains 55.2% of the variance in segmentation skills. Relative importance analysis reveals that the left AF contributes 29.53% of the variance, the left CST contributes 47.95% of the variance, and the remaining is explained by the non-significant control predictors (age and home literacy environment variables).

## DISCUSSION

In the present study, we provide for the first time evidence of prospective associations between white matter organization in infancy and subsequent language skills once children reach the school age. Until now, no structural neuroimaging study has longitudinally tracked children from infancy to school age to determine whether brain structure in infancy relates to long-term language development. Here we find relationships between white matter organization and later language skills with tract and construct specificity that account for age and key factors pertaining to the home literacy environment. Specifically, structural organization in the left arcuate fasciculus in infancy (as indicated by FA) is prospectively associated with core components of language development, phonological awareness, and vocabulary knowledge. By contrast, no prospective relationships were identified within the left inferior longitudinal fasciculus in infancy. As for the corticospinal tract, structural organization of this pathway in infancy is prospectively associated with subsequent phonological skills. These are the first results to suggest longitudinal relationships between white matter organization in infancy and language abilities in kindergarten, suggesting that an important structural foundation for language is set in infancy that underlies the subsequent developmental trajectory.

Our findings in the arcuate fasciculus build upon the literature to date implicating this pathway as crucial for language and extend beyond this literature in revealing that this tract significantly contributes to the prediction of school-age language skills from as early as infancy. Up to this point, ample evidence has established significant associations between FA in the left arcuate fasciculus and language skills among pre-school and school age children (Lebel and Beaulieu, 2009; Reynolds et al., 2019; Saygin et al., 2013). Limited previous evidence has pointed towards contributions of the arcuate from early childhood in shaping subsequent language skills, but without long-term longitudinal links from infancy due to cross-sectional and short-range longitudinal designs (Reynolds et al., 2019). Here we provide novel evidence that these previously reported links between the left arcuate fasciculus and selected language skills are not only present at the school age, but also that structural organization in the posterior portion of this tract from as early as infancy is prospectively associated with school-age language outcomes. Our findings linking the arcuate fasciculus in infancy with subsequent phonological awareness and vocabulary knowledge directly align with key constructs implicated in previous studies of school-age children (Lebel and Beaulieu, 2009; Reynolds et al., 2019; Saygin et al., 2013). These relationships also account for factors pertaining to the home environment, in line with the previously identified role of the left arcuate fasciculus in mediating the relationship between environmental exposure to language (number of conversational turns) and language skills among 4-6 year olds (Romeo et al., 2018). Our findings indicate that these relationships may be, in part, explained by pre-existing brain structure from as early as infancy, consistent with the notion that structural connectivity precedes functional development (Saygin et al., 2016). Moreover, the specificity of our findings within the posterior segment of the left arcuate fasciculus in infancy shares anatomical overlap with previous findings in preschoolers (Romeo et al., 2018). This novel evidence that structural organization of white matter in infancy is prospectively associated with language skills in kindergarten suggests that the structural foundation for the arcuate fasciculus, established within the first two years of life, contributes to the developmental trajectory of language and the corresponding brain-behavioral relationships observed in school-age children and adults.

In addition to our findings in the primary dorsal pathway implicated in language, our investigation also points towards early contributions of the corticospinal tract as a foundation for subsequent language skills. Notably, however, we did not find a comparable relationship with the ventral pathways. No significant relationships were indicated between FA in the inferior longitudinal fasciculus in infancy and language skills in kindergarten. Although the inferior longitudinal fasciculus contributes to the comprehensive network of white matter subserving language (Dick et al., 2014), our findings call for further investigation from a developmental perspective to determine which components of language may be directly associated with this tract in infancy and early childhood. By contrast, our incorporation of the corticospinal tract as a non-language-dominant tract, given its primary role in motor control, did reveal significant relationships. Here we provide the first evidence that structural organization of the left corticospinal tract in infancy is prospectively associated with phonological skills in kindergarten, specifically phonological memory and phonological awareness. Although the majority of evidence to date implicates language-dominant white matter pathways in association with phonological awareness (Lebel and Beaulieu, 2009; Reynolds et al., 2019; Saygin et al., 2013), our finding is in line with previously reported relationships between the corticospinal tract and phonological skills among 3-5 year old children (Walton et al., 2018). From a developmental perspective, the left corticospinal tract matures earlier than the arcuate and inferior longitudinal fascicles (Dubois et al., 2008; Dubois et al., 2009). Therefore, it is conceivable that higher indices of myelination within the corticospinal tract in infancy may facilitate cognitive-linguistic development in early childhood, in line with the proposition that cognitive skill development arises from domain-general interactions sub-served by multiple pathways (i.e., Interactive Specialization Theory, Johnson, 2011). Our data suggest that the corticospinal tract may play a key role from infancy in facilitating development as language-dominant tracts mature.

The present findings carry implications for language development as a process of refinement that builds upon a pre-existing structural scaffold in infancy, but also point towards a multitude of future questions. Our data support notions of the early impact of genetic susceptibility, as temporal dynamics of the transcriptome are known to be most robust during prenatal development relative to postnatal stages (Colantuoni et al., 2011; Johnson et al., 2009; Kang et al., 2011; Lambert et al., 2011; Somel et al., 2010). This is also reflected in the trajectory of white matter development, as the first two years of life signify the most rapid developmental period (Gilmore et al., 2018). While our infants were recruited within this timeframe, with an average age of 10 months it remains unclear to what extent the present findings already reflect experience-driven effects, as the first year of life encompasses a rapid time of learning and development (Dehaene-Lambertz and Spelke, 2015). Although infant age and home literacy environment at the time of neuroimaging acquisition were accounted for in the present analyses, future studies are needed that examine these relationships from a narrower and earlier starting point in infancy to disentangle biological determinants from environmental experience. For an important consideration beyond infancy, the myelination process of maturation and refinement also continues throughout early childhood into even adulthood (Barnea-Goraly et al., 2005; Lebel et al., 2008; Mukherjee et al., 2001), shaped by environmental experience and giving rise to skill acquisition (Fields, 2010). Therefore, it will be important in future work to investigate the relative contributions of white matter in infancy versus developmental changes in white matter throughout early childhood in relation to subsequent language outcomes.

One crucial consideration that remains is the respective contributions of biological determinants in the context of environmental factors that shape the developmental trajectory. Prospective associations between white matter in infancy and language skills in kindergarten in the present work are significant while accounting for factors pertaining to the home literacy environment; yet environmental factors related to parent-child interactions in the home have been previously linked with subsequent language skills in early childhood (Karrass and Braungart-Rieker, 2005; Raikes et al., 2006; Schmitt et al., 2011). Our data are to be interpreted in the context of the rich body of evidence to date implicating the important role of the environment in shaping language skills, and the modest prediction estimates of our models likely illuminate additional environmental factors that play an important role, such as the quantity and quality of exposure to language via child-directed speech from caregivers (Rowe, 2012; Weisleder and Fernald, 2013) or access to resources that facilitate education and development (Rowe, 2008). Minimal variation in parent education and income (i.e., socioeconomic status) controlled the present sample, thus, future work is necessary to determine the extent to which socioeconomic status may affect these longitudinal relationships. Nonetheless, putative influences of the home environment must be interpreted in the context of genetic influences from the parents as well, as parents contribute both environmental and genetic influences on children’s outcomes (Hart et al., 2019). Taken together, our findings illuminate the importance of investigating neurobiological foundations longitudinally from infancy in conjunction with environmental factors and potential interactions in relation to the trajectory of language development.

Overall, the present study provides the first evidence that white matter organization in infancy is prospectively associated with language abilities in kindergarten. These findings suggest that the white matter structural organization established in infancy sets one important foundation that underlies the trajectory of language development in early childhood. Our data support the working hypothesis that certain genes and their temporal dynamics, critical for brain development, establish white matter in infancy that is receptive to environmental input, and it is these factors together which ultimately shape the developmental trajectory of language acquisition. This work points towards the significance of infant brain structure in providing a scaffold upon which ongoing experience can build throughout development.

## METHODS

### Participants and Design

Forty children (20 female) were included in the present study as part of a larger NIH-funded investigation of children from infancy to school-age, tracking brain and language/preliteracy development longitudinally (NIH–NICHD R01 HD065762). Families were recruited from the Greater Boston area through the Research Participant Registry within the Division of Developmental Medicine at Boston Children’s Hospital, as well as ads and flyers disseminated in local newspapers, schools, community events, and through social media. We invited families for initial participation when children were infants (mean age at infancy: 10 months; age range: 4-18 months), and follow-up assessment was conducted when they reached the preschool/kindergarten age (mean age: 5.5 years, age range: 4 – 6.5 years). The majority of children completed follow-up assessment at the start of formal schooling (i.e., in kindergarten); three children were in first grade at the time of their follow-up. Participating children completed an MRI scan session at the infant time point and then at follow-up, a standardized battery of cognitive-linguistic assessments and repeat MRI session. Here we focus on the standardized language assessments at follow-up from the first cohort within our larger longitudinal investigation (Cohort 1 of 2 cohorts), which follows children from infancy to early elementary school. Within this cohort, 47 infants have since reached the school age and returned for follow-up assessment. Of these 47, 40 children yielded both DTI data of sufficient quality for analyses and met all criteria for inclusion at the follow-up time point, and thus were included in the present study.

Children were screened at both time points for psychiatric, neurological, or sensory illness/impairment, contraindications for MRI evaluation (i.e., metal implants), premature birth, and psychotropic medication treatment (except for one child with a diagnosis of ADHD reported at the follow-up time point). All children included were from American English-speaking families and were born at gestational week 37 or later. Of the 40 children in the present study, 29 were right-handed, eight were left-handed, and three had unknown handedness at follow-up. In addition, these children demonstrated nonverbal cognitive abilities within one standard deviation of the mean or above, as indicated by the Kaufman Brief Intelligence Test: 2nd Edition (KBIT-2, Kaufman and Kaufman, 2004). This study has been approved by the Institutional Review Board of Boston Children’s Hospital. Informed written consent was provided by a parent/legal guardian for each participating child, and written assent was provided by each child at follow-up, beginning at age four.

### Environmental Characteristics

Children were either enrolled in or had completed preschool at the time of follow-up participation. Of the 40 children in the sample, 30 participated in extracurricular activities (e.g. music, art, sports) for an average of two hours per week. There was minimal variation in socioeconomic status among families in the present sample, as indicated by parent report in a questionnaire adapted from the MacArthur Research Network (http://macses.ucsf.edu/default.php). Specifically, families were characterized by middle to high socioeconomic backgrounds, as 36 parents in the present sample had at least a Bachelor’s degree and 25 of them reported an annual income of $100,000 or more (note: one family did not provide socioeconomic information).

Another environmental factor measured included the home literacy environment (HLE), in which parents completed a questionnaire at the infant time point characterizing children’s exposure to literacy (Powers et al., 2013). Two key variables characterizing HLE were included in the present analysis: (1) the number of children’s books in the home, known to be related to children’s language and literacy-related skills (van Bergen et al., 2017), and (2) the amount of time a parent spends reading to their child, as shared parent-infant book reading has been linked with language acquisition in infancy (Karrass and Braungart-Rieker, 2005), and has been prospectively associated with subsequent language abilities and school readiness (Raikes et al., 2006).

### Language Measures

The following language assessments were investigated at the follow-up time point, which comprise a subset of the battery from the larger investigation tracking language and pre-literacy development for the purposes of the specific research questions in the present study:

#### Phonological awareness and phonological working memory

Phonological awareness was measured by two standardized subtests from the Woodcock Johnson Tests of Oral Language (WJ-IV OL, Mather et al., 2014): Segmentation and Sound Blending. The Segmentation subtest measures the ability to break words down into their basic speech sounds by requiring children to listen to words (compound, multi-syllabic, and monosyllabic) and identify the parts (at the level of compound words, syllable sounds, and individual speech sounds). The Sound Blending subtest measures the ability to synthesize speech sounds through an auditory processing task, by requiring children to listen to a sequence of syllables or speech sounds and combine the sounds into a word. Phonological working memory was measured by the Memory for Digits subtest of The Comprehensive Test of Phonological Processing (CTOPP-2, Wagner et al., 2013), in which children are required to repeat strings of numbers, which varied in length from two to eight digits.

#### Oral comprehension and vocabulary knowledge

The Oral Comprehension subtest from the WJ-IV OL was administered, which measures listening, reasoning, and vocabulary abilities by requiring children to listen to a brief audio-recorded passage and then provide the omitted word through the use of syntactical and semantic clues. The Peabody Picture Vocabulary Test (PPVT-4, Dunn, 2007) measures vocabulary knowledge, in which the examiner reads a single word and children are asked to select the picture (out of four possible choices) that best reflects that word.

These assessments were administered and scored by a trained psychoeducational evaluator, and then double-scored by an additional evaluator to ensure accuracy. Raw scores for each of the assessments were converted to standard scores, and standard scores were utilized in subsequent analyses.

### Neuroimaging Acquisition

Neuroimaging was acquired while the infants were naturally sleeping, utilizing a pediatric neuroimaging protocol previously established (Raschle et al., 2012). Diffusion-weighted and structural T1-weighted images were acquired on a 3.0 Tesla Siemens TrioTim MRI scanner with a standard Siemens 32-channel radio frequency head coil. One parent remained in the MRI room with the infant for the duration of the scan, in addition to a researcher who stood by the bore to monitor changes in the infant’s sleeping state and potential motion. Structural T1-weighted whole-brain multi-echo magnetization-prepared rapid gradient-echo sequences with prospective motion correction (mocoMEMPRAGE) were acquired for each participant (acquisition parameters: TR = 2270 ms; TE = 1450 ms; TA = 4.51 min; flip angle = 7°; field of view = 220 x 220 mm; voxel resolution = 1.1 x 1.1 x 1.0 mm (176 slices) with an inplane acceleration factor of 2). Diffusion-weighted echo planar images were acquired using 64 slices from 30 gradient directions with 10 non-diffusion-weighted volumes (acquisition parameters: slice thickness = 2.0 mm; b = 1000 s/mm^2^; field of view = 256 x 256 mm; TE = 88 ms; TR = 8320 ms; TA = 5:59 min; flip angle = 90°).

### Diffusion-Weighted Image Processing

Diffusion-weighted images were processed with the approach taken in a previous study from our lab with infants from this cohort(Langer et al., 2015). A brain mask was generated from the structural T1-weighted image utilizing the Brain Extraction Tool (BET, Smith, 2002) to separate brain tissue from nonbrain tissue. Raw DWI data were converted from DICOM to NRRD format through the DicomtoNRRDConverter software of Slicer4 (www.slicer.org). Quality control of diffusion images was evaluated utilizing QCTool, a MATLAB-based toolbox that detects artifacts based on Kullback–Leibler divergence calculations for each gradient in the sequence relative to a brain intensity mask for each individual subject. A combination of the tool’s recommendations and visual inspection was implemented to detect motion artifacts, and gradients containing motion artifacts were removed prior to tract estimation. Following quality assurance, DWI data were processed with the VISTALab mrDiffusion toolbox and diffusion MRI software suite (www.vistalab.com), including eddy current correction and tensor-fitting estimations with a linear least-squares (LS) fit for fitting the diffusion tensors.

### Automated Fiber Quantification

The Automated Fiber Quantification (AFQ) software package (github.com/jyeatman/AFQ, Yeatman et al., 2012b) was used to quantify specific white matter tracts in the current study, in accordance with previous procedures and parameters employed in our laboratory (Langer et al., 2015). For a brief summary, whole brain tractography was computed using a deterministic streamline tracking algorithm (Basser et al., 1994; Mori et al., 1999), with an FA threshold of 0.1 (as previously employed with this age range, Langer et al., 2015). Fiber tracking was terminated in instances where the minimum angle between the last path segment and the next step direction was > 40°. Region of interest (ROI)-based fiber tract segmentation and fiber-tract cleaning were then employed (using a statistical outlier rejection algorithm) before FA quantification was conducted along the trajectory of each tract based on eigenvalues from the diffusion tensor estimation (Basser et al., 1994). FA of each fiber was sampled to 100 equidistant nodes. For each tract, the characterization of 100 nodes was resampled to 50 nodes, thereby discarding the portion of the tract where individual fibers separate from the core fascicle toward their destination in the cortex. This method was employed based on previous literature (Langer et al., 2015; Wang et al., 2016), since it is known to improve normalization and co-registration of each tract for group comparisons (Yeatman et al., 2011). For the AF, resampling of nodes was furthermore employed utilizing an alignment approach previously established in the AFQ literature (Langer et al., 2015; Yeatman et al., 2012a). Specifically, the trajectories of FA among all participants were aligned to the lowest point of FA along the trajectory of each tract to co-register anatomically analogous nodes of the tract across subjects and ensure comparison of comparable regions in the principal arc of the tract, in accordance with the previously established protocol (Yeatman et al., 2012a).

Since AFQ utilizes a conservative, automated approach for tract identification, tracts of interest could not be identified in all infants. Automated tractography of the ILF and CST was successful for all infants, and the left AF was successful for 34 out of 40 infants. Although we initially set out to also investigate the superior segment of the arcuate, the superior longitudinal fasciculus, we were unable to reliably reconstruct the tract utilizing this automated approach in a sufficient number of infants to warrant inclusion in the present analysis. Therefore, the present analysis investigated the left-hemispheric AF and ILF, with a focus on the left hemisphere given the well-established left-hemispheric dominance for language in previous literature (Catani et al., 2007; Eluvathingal et al., 2007; Glasser and Rilling, 2008; Lebel and Beaulieu, 2009; Parker et al., 2005; Powell et al., 2006). We also incorporated the left corticospinal tract to examine whether potential effects are specific to language-related tracts or extend beyond within the primary motor tract.

### Statistical Analyses

Prospective associations were examined by employing multiple regression analyses with the FA of left-hemispheric white matter pathways in infancy as predictors, key characteristics of the home literacy environment in infancy (number of children’s books in the home, amount of time spent reading with the child at home) as control variables, and subsequent pre-school age measures of language as outcome variables. These relationships were examined controlling for infant age in order to account for variability in age at time of scan in infancy. Multiple regression analyses were conducted in MATLAB (https://www.mathworks.com/products/matlab.html) and R (https://www.r-project.org/).

In addition to the classical procedure of correcting for multiple comparisons, parametric bootstrapping was adopted to simulate the sampling distribution in the general population in order to ensure proper evaluation of the magnitude of the present effects. This approach involved simulating the population distribution based on partial beta values using sampling with replacement from the original sample. A simulation of 5,000 replicated samples of the same size was programmed in MATLAB and replicated in Mplus version 8, using variance estimates of the correlation coefficient up to 8 decimal places and then simulating the population distribution using mean and variance estimates of the sample’s correlation coefficient. The distributions were tested for normality using the Kolmogorov-Smirnov test and across all bivariate comparisons they did not deviate from normality besides the expected nominal alpha level; thus symmetric 95% confidence intervals were utilized. Final indicators of significance for all analyses models were derived from this parametric bootstrapping procedure. In a final step, correction for multiple comparisons was employed utilizing the Family-Wise Error (FWE) correction method.

## Acknowledgements

This work was funded by NIH–NICHD R01 HD065762, the William Hearst Fund (Harvard University), and the Harvard Catalyst/NIH (5UL1RR025758) to N.G. and P.E.G., and the Sackler Scholar Programme in Psychobiology to J.Z. We would like to thank all participating families for their long-term dedication to this study. We are grateful for all additional members of the research team who contributed to data collection and quality control, especially Emma Boyd, Lilla Zollei, Bryce Becker, Danielle Silva, Sara Smith, Ted Turesky, and Doroteja Rubez. We also thank Carolyn King for her feedback on the manuscript.

## Author Contributions

N.G. and P.E.G. designed the study. B.G. oversaw and implemented MR sequences. J.Z., J.S., M.F., J.D., and C.C. collected the data. J.Z., X.Y., G.S., and N.G. analyzed the data with assistance from J.S., M.F., J.D., and C.C. J.Z. and N.G. wrote the manuscript; coauthors provided feedback and edits on the manuscript.

## Declaration of Interests

The authors declare no competing interests.

